# Detecting m6A RNA modification from nanopore sequencing using a semi-supervised learning framework

**DOI:** 10.1101/2024.01.06.574484

**Authors:** Haotian Teng, Marcus Stoiber, Ziv Bar-Joseph, Carl Kingsford

## Abstract

Direct nanopore-based RNA sequencing can be used to detect post-transcriptional base modifications, such as m6A methylation, based on the electric current signals produced by the distinct chemical structures of modified bases. A key challenge is the scarcity of adequate training data with known methylation modifications. We present Xron, a hybrid encoder-decoder framework that delivers a direct methylation-distinguishing basecaller by training on synthetic RNA data and immunoprecipitation-based experimental data in two steps. First, we generate data with more diverse modification combinations through in silico cross-linking. Second, we use this dataset to train an end-to-end neural network basecaller followed by fine-tuning on immunoprecipitation-based experimental data with label-smoothing. The trained neural network basecaller outperforms existing methylation detection methods on both read-level and site-level prediction scores. Xron is a standalone, end-to-end m6A-distinguishing basecaller capable of detecting methylated bases directly from raw sequencing signals, enabling de novo methylome assembly.

---

RNA modification plays essential roles in various biological processes, including stem cell differentiation and renewal, brain functions, immunity, aging, and cancer progression [1–4]. Among the various types of RNA modifications, N6-Methyladenosine (m6A) is one of the most abundant versions and is involved in various biological processes including mRNA expression, splicing, nuclear exporting, translation efficiency, RNA stability, and miRNA processing [1]. Accurate detection and quantification of m6A modifications is crucial for understanding their impact on gene regulation and cellular processes [5, 6].

Next-generation sequencing (NGS) technologies identify nucleotides through a synthesis process, leading to the loss of post-transcriptional information [7]. Therefore, indirect methods are required to detect RNA modifications with NGS. These approaches first isolate the modified RNA and then conduct reverse transcription and cDNA sequencing to reveal the modifications. Two primary strategies are used to experimentally isolate RNA modifications. One type of approach involves immunoprecipitation. Examples of methods using this approach include MeRIP-Seq [8], m6A-Seq [9], PA-m6A-Seq [10], m6A-CLIP/IP [11], miCLIP [12], m6A-LAIC-Seq [13], m6ACE-Seq [14], and m6A-Seq2 [15]. These methods rely on antibodies that target the modified ribonucleotide and enrich the RNA fragments with the target modified bases. The other type of approach is chemical-based detection. Examples of methods using this approach are Pseudo-Seq [16], AlkAniline-Seq [17], Mazter-Seq [18], m6A-REF-Seq [19], DART-Seq [20], RBS-Seq [21], and m6A-SAC-seq [22]. These techniques use chemical compounds or enzymes that selectively interact with the modified ribonucleotide, either cleaving or modifying the RNA reads to halt or disturb the reverse transcription process. This is followed by short-read cDNA sequencing, which identifies the RNA modifications by comparing the read ends of the cDNA or the base mismatches/deletions in cDNA. Although these methods were able to generate detailed maps of RNA modification sites, they all use external compounds which makes it hard to obtain the required single base resolution. They also face other challenges and shortcomings including the limited availability of antibodies or compounds for specific modifications [23], nonspecific antibody binding [24–26], low single-nucleotide resolutions [8, 9], and, importantly, an inability to identify the exact location of a modification.

Direct RNA sequencing using nanopores offers a promising alternative [27]. An RNA molecule can be sequenced by measuring the intensity of the current flowing through the pore as the RNA molecules pass through it. Modified RNA nucleotides produce different signals than their unmodified counterparts, providing information about the modifications at the single-molecule read resolution [28, 29]. However, to detect specific modifications from subtle signal changes we need an optimized algorithm, which is normally obtained through supervised learning or a comparative approach [30]. Unfortunately, current data are not immediately suitable for supervised learning due to the lack of experimental techniques for identifying the methylation state at the single-read resolution.

*In vitro* transcription (IVT) data, which are transcribed from either experimentally synthesized DNA sequences or native DNA [28, 31], can provide reads that are either completely methylated or not methylated at all (all-or-none), but the diversity of the sequence compositions in synthesized DNA datasets is limited due to the constraints concerning the maximum DNA length that can be synthesized and the associated costs. In addition, the IVT dataset lacks partially methylated reads with known methylation states. Although partially methylated reads can be generated by introducing a mixture of modified and canonical adenine during *in vitro* transcription, the location of methylation remains unknown because in such mixtures the RNA polymerase randomly selects adenine from either type during the transcription process. Models trained to identify modifications on all-or-none modified reads perform poorly on biological reads, which are usually sparsely methylated [31, 32]. Methods using such synthesized datasets include training a classifier to predict sequence segments (5-mers) given their corresponding nanopore raw signal segments [33] or features of these segments [28, 29, 31, 34]. The signal segments are extracted from raw signal after performing base-calling and alignment, using models trained on canonical data (data with no methylation). As we show, the performance of such a classifier is limited since it is only trained on isolated short segments, losing contextual information. In addition, these models are trained solely on manually selected features including mean, standard deviation, and duration of isolated signal segments corresponding to 5 bases, which can lead to the loss of more detailed signal information. Recently, a new method, CHEUI, was trained using longer signal segments, yielding impressive results on IVT data [35]. However, it suffers from overfitting when applied to real biological samples (Fig. 2, [36]).

Immunoprecipitation (IP) data from assays such as m6ACE-seq and m6A-CLIP-seq relies on the use of antibodies [11, 12, 37]. However, this strategy works at a high level. It only provides the modification proportion for each reference transcriptomic position, i.e., a site-level modification rather than the modification state for each individual read (read-level). m6Anet [36] employs multiple-instance learning [38] to train a classifier using IP data leading to improved site-level accuracy. However, IP data misses many methylation sites, particularly in low-coverage regions [25]. Additionally, due to nonspecific antibody binding, the methylation detection results obtained through immunoprecipitation experiments produced a false-positive rate of approximately 11%, which can vary between studies [18, 39]. M6Anet also requires a minimum coverage level of 20 reads for a site to be detected due to the way the model is trained. The training involves maximizing the probability of detecting at least one methylated read among the reads covering a known methylated site. Such coverage depth is not always available. Finally, as in the other existing models, m6Anet relies on a basecaller and segmentation tools that are trained on nonmodified reads (canonical reads).

In summary, previous approaches try to identify m6A sites using basecalling errors [28, 29, 31, 34], by comparing between control samples [29, 40], trained on IVT data [33, 35] or trained on noisy labels from IP data [36]; these methods are summarized in Tables 1 and S1. As we will show, the fact that they are only trained on one type of data limits their performance (Figure. 2a,b and Supplementary Figure. 3).

**Table 1.**
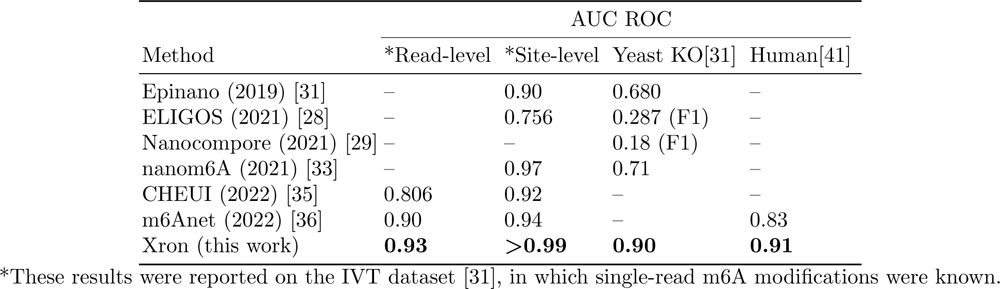
Reported Performance of m6A Modification Identification Achieved by Existing Works.

We present a method that takes a different approach by detecting methylation during the basecalling phase. We predict methylated bases directly from the long current signal by training a methylation-distinguishing basecaller. To achieve this, we developed Xron, a hybrid encoder-decoder framework (Fig. 1). The encoder is a convolutional recurrent neural network (CRNN) encoding the observable signal into a kmer representation. After it has been trained and fine-tuned, the CRNN serves as a methylation-distinguishing basecaller for new data. The decoder is a nonhomogeneous hidden Markov model (NHMM), which serves as a generative model for achieving signal segmentation and alignment when preparing the training dataset. Applying the NHMM, we created a partially methylated dataset to train the CRNN and produce a methylation-distinguishing basecaller. The CRNN is then fine-tuned using the IP data, further enhancing the basecaller’s generalizability. This framework enables us to obtain a highly accurate methylation-distinguishing basecaller by exploiting both IVT data and IP data. This approach outperforms all previous methods on synthesized and biological samples and provides a comprehensive, end-to-end solution for methylation base detection.

**Fig. 1.**
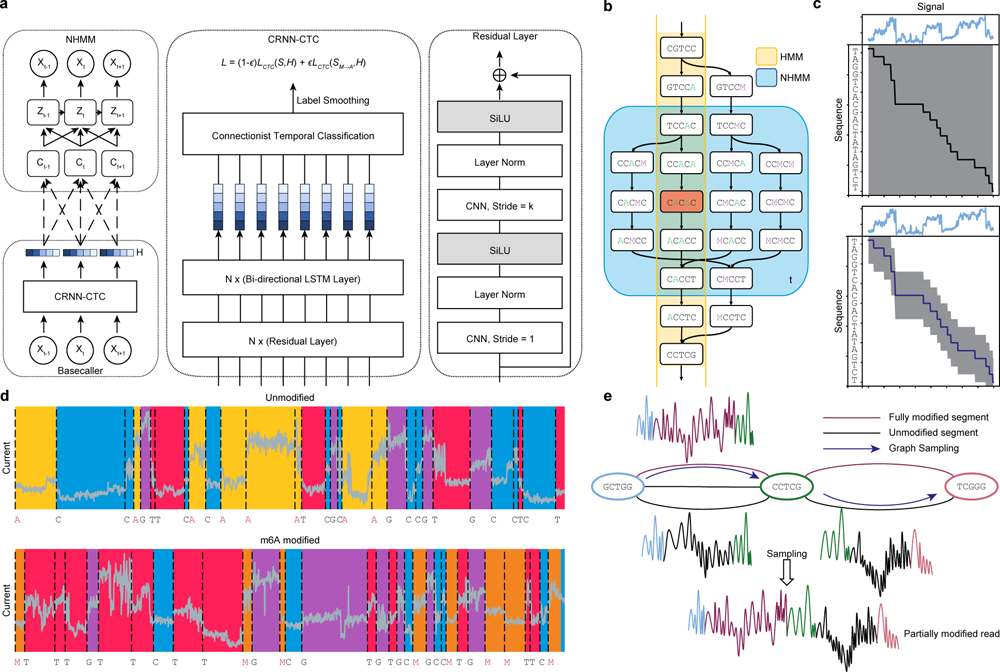
Schematics of Xron model and the data augmentation process through crosslinking and sampling. **a**, Xron consists of two parts: a nonhomogeneous hidden Markov model (NHMM) and a convolutional recurrent neural network (CRNN) with a connectionist temporal classification (CTC) decoder. **b**, Comparison between HMM and NHMM. The transition matrix of a HMM (yellow) encodes the whole Markov chain of k-mers, while the transition matrix of the NHMM (blue) at time *t* only encodes the Markov chain of the five nearby k-mers given the predicted k-mer (shown in red) at time *t*. The Markov chain is also expanded to include the k-mers with all combinations of the A and M (m6A) bases. We create partially methylated reads using data augmentation, first segmenting the signal and then cross-linking the reads and their corresponding signal in silico. To achieve this, we design a novel nonhomogeneous hidden Markov model (NHMM) that can be trained to conduct signal segmentation in a semi-supervised fashion on modified reads, even when lacking methylation labels. The NHMM is trained using the forward-backward algorithm with its transition matrix conditioned on a canonical baseaclled sequence and its alignment, thus giving the maximum likelihood estimation of the model parameters regarding methylation base. The Viterbi path of the NHMM gives the alignment between the current signal and sequence. Following the signal segmentation process performed with the NHMM, the NHMM was used to create a training dataset with partially methylated reads and their true labels for methylation detection training by augmenting all-or-none modified reads. **c**, The transition process of the NHMM is constrained by the neural network’s output, leading to a smaller probability space and making it easier for the model to find the optimal alignment. **d**, The NHMM is trained in a semi-supervised manner on IVT datasets, including fully modified, unmodified, and partially modified reads. It provides accurate signal segmentation results for both unmodified and modified sequences. **e**, In-silico read crosslinking. The fully modified or unmodified reads are first broken into segments at the invariant k-mers to form a signal-k-mer graph, whose nodes are k-mers and whose edges are signal segments. Then, a partially methylated read is sampled from the k-mer signal graph.

**Fig. 2.**
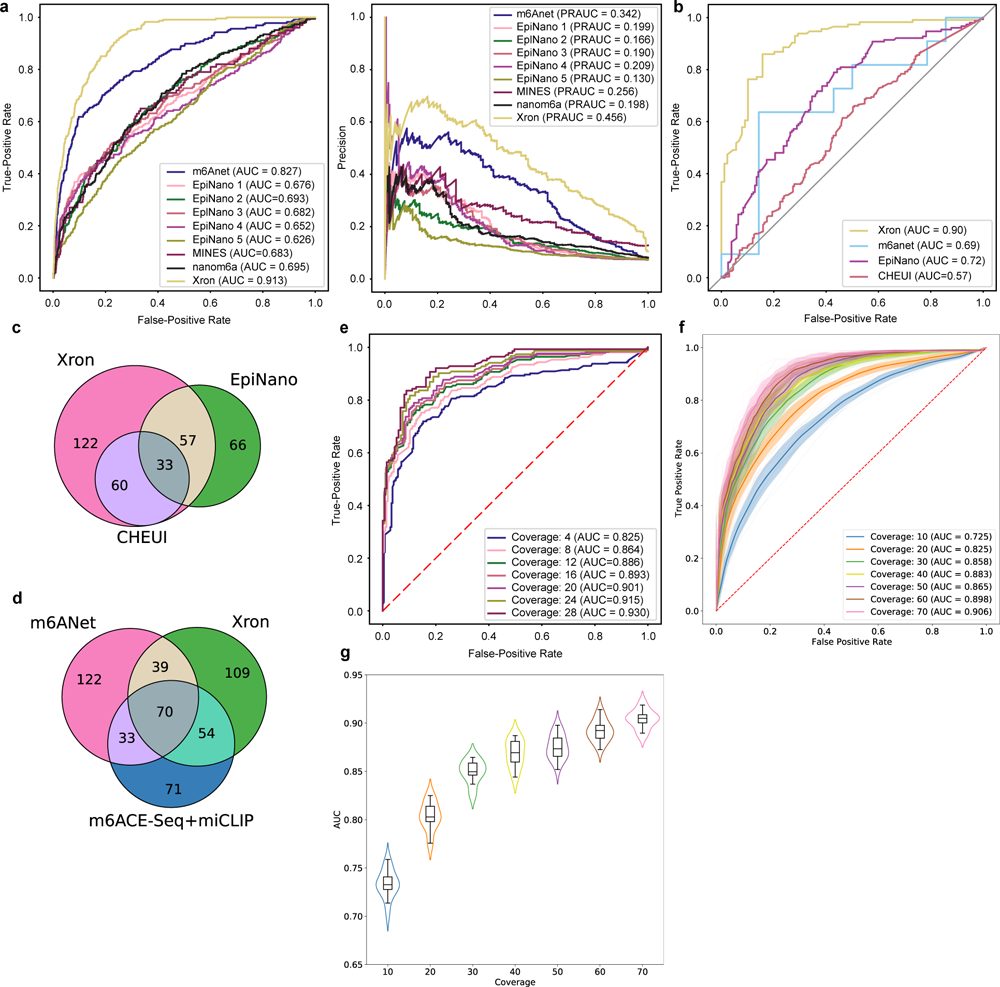
Comparison of Xron models across two different species. **a**, ROC and PR curves of m6A prediction on human HEK293T cell line, produced by Xron and other models. **b**, ROC curves produced by Xron and other models on yeast data. **c**,**d**, Venn diagram showing the overlapping sites predicted by Xron and other methods on Yeast (**c**) and HEK293T (**d**) data. **e**, ROC curves produced by Xron for detecting m6A methylation in yeast data under different minimum sequence coverage thresholds. **f**, ROC curves generated by Xron for detecting m6A methylation in down-sampled yeast data with different coverage. **g**, Distribution of AUC score of Xron on down-sampled yeast data.

## Results

### Applying Xron to identify m6A methylation on direct RNA sequencing datasets

Xron performs methylation-distinguishing basecalling, outputting methylated bases directly from the raw sequencing signal emitted from the nanopore. Its neural network basecaller is trained on an augmented partially methylated dataset and then fine-tuned using IP data. We tested Xron on three public direct RNA sequencing datasets: an IVT dataset [31], a yeast dataset [31], and a human embryonic kidney cells (HEK293T) dataset [36].

The IVT dataset [31] was synthesized from artificially designed sequences followed by *in vitro* transcription. The dataset contains either fully methylated or fully unmethylated reads. Signal intensity shows differences around the center base of the kmer between modified and unmodified sites (Fig. 3a and Supplementary Fig. 1). The sequences are designed to contain all 5-mers, including the most common k-mer (GGACT) and all 18 DRACH motifs (Fig. 3a,b).

**Fig. 3.**
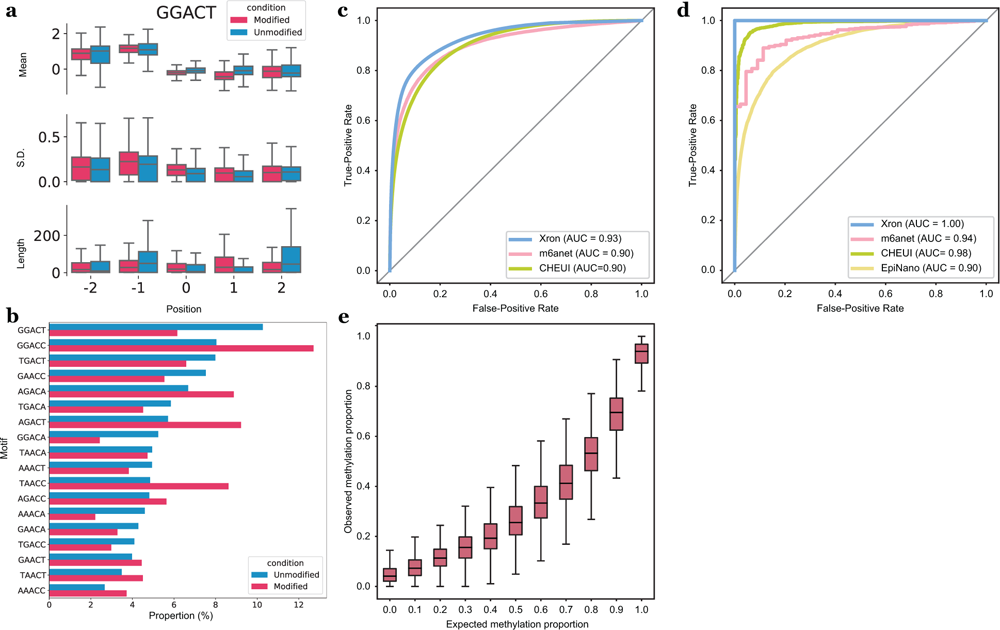
Evaluation of the m6A detection results obtained for synthesized IVT RNA reads and stoi-chiometry prediction. **a**, Box plot comparing the distribution of the mean, standard deviation, and length for the signal segmented by NHMM with 5, 232 modified sites and 18, 464 unmodified sites for the GGACT motif. Horizontal lines show the median, the box denotes the interquartile range, and the whiskers extend to 1.5 times the interquartile range. Points beyond this range are considered outliers and are removed from the plot. **b**,**c**, ROC curves of Xron against m6anet and CHEUI for read-level (**b**) and site-level (**c**) m6A modification predictions. **d**, Bar plot showing the relative proportion of DRACH 5-mer motif for 84, 919 modified and 179, 717 unmodified positions. **e**, Box plot showing the m6A ratio predicted by Xron with different proportions of IVT control and IVT m6A RNA mixing.

The yeast dataset [31] contains direct RNA sequencing reads from two strains, a wild-type strain, and a “ime4Δ” knockout strain, in which IME4 was deleted. The deletion of IME4 results in the complete elimination of m6A bases, making it a negative control. The yeast dataset contains three independent biological replicates for each strain. Two were used in this study; the first replicate was used for training, and the second was used for evaluation.

The human HEK293T cell dataset [36] contains direct RNA-Seq data from the HEK293T cell line [34], with methylation sites identified by m6ACE-Seq [14] and miCLIP data [12] on the same cell line. The dataset contains three replicates, and we used the first replicate to evaluate the method. (See Methods for details about replicates and datasets used for training and evaluation.)

### Xron accurately identifies m6A sites

To evaluate the performance of Xron, we applied Xron that is finetuned on yeast data to direct RNA sequencing data derived from the human HEK293T cell line [34]. Although Xron is pre-trained using human IVT reads (Methods), no human methylation information is used during training since all human reads are canonical. To validate the model, we used the m6A sites detected by m6ACE-Seq and miCLIP from the human HEK293T cell line as the true labels during evaluation, following previous work [36]. We used the m6A sites identified by m6ACE-Seq and miCLIP as positive samples and the other sites with the same 5-mer as negative samples. Xron achieved the best ROC AUC of 0.91 (Fig. 2a) compared with those of Epinano (0.69) and m6Anet (0.83) and the best precision-recall (PR) AUC of 0.456 (Fig. 2a) compared to m6Anet (0.342) and MINES (0.256).

### Xron is sensitive to IME4 knockouts

In addition, we also evaluated Xron on a yeast dataset using a ime4Δ knockout *S. cerevisiae* strain where the m6A modification was completely eliminated [37] as the control dataset, following a previous study [31]. We used the second replicate sample of the dataset for evaluation, as we had fine-tuned Xron on a subset of the first replicate. We treated the m6A sites in the wild-type strain as modified sites and the same sites in the ime4Δ knockout strain as unmodified sites. We compared Xron with other models for predicting modified/unmodified sites. Xron achieved an AUC-ROC score of 0.90 (Fig. 2b) on this task, providing a 21% increase over the second-best model, Epinano (0.72).

### Xron detects more methylation sites and achieves high accuracy under low coverage settings

As m6anet intrinsically requires a minimum coverage of at least 20 to obtain site methylation predictions, this results in a much smaller sample size (11 sites detected). In the same setting, Xron yields 171 sites with a minimum coverage of 20 on the yeast dataset, which results in higher AUC-ROC accuracy than m6anet (0.90 versus 0.69). In total, Xron detects 272 sites reported in the IP data, compared to the 156 sites detected by Epinano and the 93 sites detected by CHEUI (Fig. 2c). Sites detected by Xron also show higher support from the IP technique (124) compared to m6Anet (107) in the HEK293T cell line (Fig. 2d). While different methods identify various m6A methylation sites, many sites are detected exclusively by one method. This observation aligns with previous reports [14, 36]. We next tested if including more low-coverage sites by setting different minimum sequencing coverage thresholds would influence the prediction accuracy of Xron (Fig. 2e). We found that increasing the read coverage yielded superior site-level methylation prediction accuracy, increasing from a 0.825 AUC-ROC score for a minimum read coverage level of 4 to a 0.930 AUCROC score with a minimum read coverage level of 28. This suggests that with higher sequencing depth, Xron can further enhance the precision and accuracy of methylation detection. Meanwhile, Xron outperforms other models by a large margin even when setting the minimum read coverage level to 4, with AUC 14% more than the second best model, Epinano (0.825 versus 0.72). Furthermore, to evaluate Xron’s performance in low-coverage regions, we down-sampled the reads to limit the maximum coverage at each site to a range of 10 to 70. Xron achieved an accuracy of 0.725 with maximum coverage of 10, outperforming other models with full data (Fig. 2f,g).

### Xron achieves nearly optimal site-level prediction on a synthesized RNA dataset

We evaluated Xron on a synthesized RNA IVT dataset [31] obtained from a different replicate than the training dataset (see the Methods section). In this dataset, the true methylation modifications were known for each position in each read, as the reads were either from a fully modified or a fully unmodified run. Our model achieved an AUC ROC of 0.93 on the single-read-level prediction task (Fig. 3c), in which the model has to predict m6A bases or A bases for each read at RRACH sites identified by previous antibody immuno-precipitation experiments [37]. Our model outperforms the second-best read-level model (m6anet) by 3% (0.93 versus 0.90) and an almost optimal AUC ROC of >0.99 for site-level prediction (Fig. 3d), outperforming the second-best site-level model (CHEUI) by nearly 2% (≈ 1 versus 0.98).

### Xron provides m6A stoichiometry

By aligning the reads to the reference genome and piling up the single-read m6A modification predictions for different sites, Xron can predict site-level m6A modification stoichiometry, i.e., the fraction of modified bases at a site. We evaluated this ability using a synthetic dataset.

The dataset was a mixture created by randomly sampling reads from fully modified or unmodified IVT datasets [31] with specific mixture proportions, which included 0%, 10%, 20%, 30%, 40%, 50%, 60%, 70%, 80%, 90%, and 100%. We calculated the model-predicted m6A proportion as the number of m6A bases called per site divided by the total number of reads aligned to this site. The median relative modification proportion followed the same trend as the expected methylation proportion. The trend in stoichiometry level was successfully recovered (Fig. 3e).

### Xron performs consistent basecalling on m6A-modified datasets

To compare the performance of Xron as a basecaller with a canonical basecaller, we evaluated the basecalling accuracy of Xron and compared it with that of the Guppy ONT basecaller (Table 2 and Supplementary Table S2). We evaluated the basecall quality achieved on three datasets: the synthesized IVT RNA dataset, the S. cerevisiae yeast dataset, and the human HEK293T cell line dataset, considering both modified and unmodified reads. For the synthesized IVT RNA and yeast datasets, we used the second replicate, which was not used as training data. Xron suffers less (or no) accuracy drop on datasets with m6A modifications. It exhibited no performance loss on datasets with methylation compared to the control dataset. On the other hand, Guppy showed performance decreases on all three datasets with methylation compared to its performance on the unmodified control datasets, including a 14.47% drop in the identity rate on the synthesized reads and a 7.55% drop in the identity rate on the HEK293T reads.

**Table 2.**
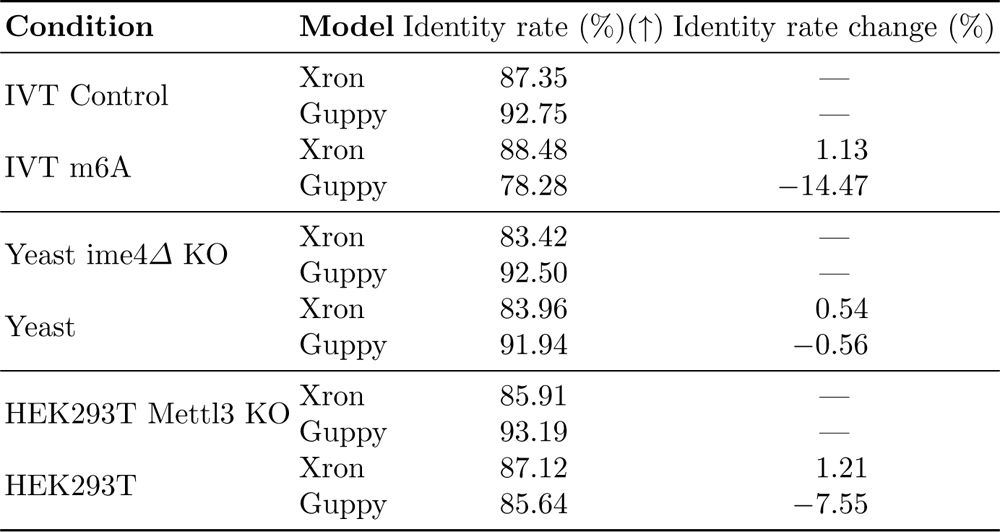
Accuracy comparison between Xron and Guppy on three different datasets and their control datasets. The identity rate (%) was defined as the number of matched bases in the query sequence divided by the number of bases in the reference sequence (the higher the better). All reported rates are mean values among the aligned reads.

## Discussion

Several computational methods [28, 29, 31, 33, 35] have been used to detect m6A methylation. These methods require accurate training data, usually obtained using synthesized RNA reads containing the modification of interest, obtained through experimental methods such as m6ACE-Seq or miCLIP, or from a comparative analysis against control data. However, these methods exhibit a performance drop when they are applied to other datasets, implying the existence of overfitting. In addition, these methods usually can only provide site-level methylation, losing read-level resolution. We developed an end-to-end m6A modification detection system for nanopore direct RNA sequencing and, for the first time, created an m6A-distinguishing base caller. Our system, Xron, includes an NHMM model for k-mer decoding and a neural network basecaller. By employing data augmentation and semi-supervised learning, we constructed an NHMM that is capable of performing accurate signal sequence alignment and introduced a novel training dataset for m6A methylation detection. The training pipeline established in our work facilitates supervised basecaller training without necessitating complex feature engineering and using both IVT and IP data available to overcome overfitting.

Quantifying the transcriptome-wide modification rates is one of the key challenges in methylation detection. From the read-level methylation states given by Xron, the modification stoichiometry for each site can be obtained. Meanwhile, our method does not require a high minimum coverage depth, which is essential for detecting methylation in low-expression regions. Comparative methods detect methylation by analyzing data from different conditions [29, 34]. While Xron does not require a control sample to detect methylation, it can also facilitate the use of a control sample by comparing the same site across samples. In addition, compared to other methods where the model performance is influenced by aspects such as base-calling algorithms, accuracy in the alignment of the reference sequence to signal, and segmentation of the raw signal, Xron reads out methylation information directly from the raw signal. More training data on different experimental protocols and different organisms will likely further improve the accuracy of Xron and other supervised approaches, while the training framework of Xron can easily adopt these additional training data into the finetuning pipeline.

As a basecaller, Xron achieves a consistent identity rate among methylation and unmethylation datasets. Although there is a performance gap in terms of identity rate between Xron and the basecaller Guppy, this is likely due to the different neural network architecture used. In future research, it would be beneficial to investigate various neural network structures since previous studies have shown that alterations to the convolutional-recurrent neural network architecture can yield enhanced basecalling accuracy. For example, Guppy uses QuartzNet [42], a neural network designed initially for speech recognition. SACall [43] employs an attention mechanism, while RODAN [44] integrated squeeze-and-excitation [45] layers into a base CNN.

Currently, the NHMM takes only raw signal as its input. This has several advantages, including being easy to train and having a closed-form solution for parameter estimation. However, additional input features can be added to the NHMM, including the encoded representation from the neural network base caller.

Xron was used to detect m6A modification, however, our framework is suitable for training a basecaller for detecting any natural post-transcription modification, including DNA methylation such as m5C and other types of RNA modification. Xron can also be retrained to detect artificial modifications at a single-molecule level, such as detecting modifications introduced in small non-coding RNA [46].

## Acknowledgements

This work was supported in part by the US National Science Foundation [DBI-1937540, III-2232121], the US National Institutes of Health [R01HG012470], and by the generosity of Eric and Wendy Schmidt by recommendation of the Schmidt Futures program. We also thank the Pittsburgh Supercomputing Center for providing computational resources through the Bridges2 system. H.T. is supported by funding from Oxford Nanopore Technologies plc and the School of Computer Science, Carnegie Mellon University - The Joint CMU-Pitt Ph.D. Program in Computational Biology (CPCB). Conflict of Interest: C.K. is a co-founder of Ocean Genomics, Inc. H.T. is supported by funding from Oxford Nanopore Technologies plc. M.S. is an employer of Oxford Nanopore Technologies plc. We thank Minh Hoang for reviewing the manuscript and offering valuable feedback. We thank Tim Massingham (XGenomes Corp.) for the helpful discussion on signal segmentation.

## Code Availability

Code is hosted at GitHub repository https://github.com/haotianteng/xron. Xron is available under a GNU GENERAL PUBLIC LICENSE v3.0. Xron is built with Python 3.8 and PyTorch 1.12, and has been tested on PyTorch 1.13 and 2.0. We used ChatGPT to correct grammatical errors and improve the flow of early drafts of this manuscript.

## Data Availability

The IVT RNA datasets were obtained from Epinano project [31] through the GEO database (GSE124309). The ELIGOS IVT RNA datasets were obtained from ELIGOS project [28] through the SRA database (SRP166020). The Yeast datasets (wild and ime4-knockout) were obtained from Epinano Project [31] through the GEO database (GSE126213). The HEK293T cell lines data were obtained from the SG-NEx Project [41] through ENA (PRJEB40872).

## Methods

Xron is trained using both IVT and IP datasets to obtain better performance. It was first trained on a surrogated IVT dataset and then fine-tuned on IP data. To make efficient finetuning and to avoid overfitting to the all-or-none methylated reads in IVT data when training with the long current signal, we create partially methylated reads using data augmentation, first segmenting the signal and then cross-linking the reads and its corresponding signal in silico. To achieve this, we design a novel nonhomogeneous hidden Markov model (NHMM) that can be trained to conduct signal segmentation in a semi-supervised fashion on modified reads, even when lacking methylation labels. The NHMM is trained using the forward-backward algorithm with its transition matrix conditioned on a canonical basecalled sequence and its alignment, thus giving the maximum a posteriori estimation of the model parameters regarding methylation base. The Viterbi path of the NHMM gives the alignment between the current signal and sequence. Following the signal segmentation process with the NHMM, we prepared a partially methylated dataset through data augmentation, splicing the fully methylated and unmethylated segments. Training on this augmented dataset diminishes the inductive bias of the model on partially methylated reads when training with entirely methylated or nonmethylated reads. We then trained an end-to-end methylation-detection basecaller on the augmented dataset, and it achieved high-accuracy methylation base detection at a single-read resolution. We further improved the basecaller by applying a fine-tuning procedure on IP data with label smoothing to obtain a more accurate basecalling model. Finally, we benchmarked different m6A detection methods on three datasets, including a synthetic IVT dataset, a yeast dataset, and a human HEK293T cell line, demonstrating that Xron yields accurate methylation-aware basecall and generalizes to different species.

### NHMM trained using semisupervised learning

We design a hybrid framework to conduct signal segmentation and alignment when methylated bases are present. A homogeneous HMM (we refer to this model as an HMM throughout the remainder of this paper for convenience), as employed in the common Nanopolish preprocessing tool [47], faces challenges when applied to sequences with methylation bases. The absence of ground truth for the methylation states in each basecalled sequence prevents supervised HMM training. However, training the HMM unsupervised, using only signal and reference genome, is difficult due to the high noise contained in nanopore sequencing signals, the long lengths of the electrical signals, and the similar signal levels between certain k-mers and their methylated counterparts. Additionally, totally unsupervised training is not necessary as we already have the canonical basecalled sequence with alignment given by the canonical basecaller and the reference genome. Although the signals are error-prone in the methylated region, they still provide a general sketch of the sequence. Thus, instead of performing unsupervised learning with the HMM, we develop a semi-supervised training process using an NHMM, where we use the basecalled canonical sequence as a prior when building the transition chain backbone in the NHMM. In contrast with an HMM possessing a homogeneous transition matrix that remains constant over time t, an NHMM possesses a nonhomogeneous transition matrix that depends on the external variables and varies over time t, allowing the use of dynamic control for the transition process. Various NHMMs have been used in meteorology [48] and economics [49, 50] by constructing transition matrices that depend on time-varying covariates, such as seasonality [48] or economic cycle indicators [50]. In our case, the base probabilities along time t predicted by an existing canonical basecaller (a base caller trained to predict only canonical bases) are used as the time covariates of the transition matrix. This approach enables the model to concentrate on the section of the Markov chain guided by the predicted base probability (Fig. 1c), rather than dealing with the entire chain as is required in unsupervised learning using HMM, which is more challenging and error-prone.

### NHMM for methylated sequence segmentation and alignment

The NHMM represents the input sequence of raw current signals as X = (x_1_, . . ., x*_T_*) for a given k-mer sequence Z = (z_1_, . . ., z*_T_*) inside a nanopore over the sequencing duration T. Each signal point x*_t_* represents a normalized current value, while z*_t_* is a variable indicating the k-mer at time t. The transition matrix of the NHMM is constrained on the basecalled sequence and its alignment given by the canonical basecaller. More specifically, suppose we are given the base probability matrix H = (h_1_, . . ., h*_T_*) ∈ R*^B^*^×*T*^, where B is the number of bases and h*^b^* is the probability of base b at time t, which is obtained from an existing canonical neural network basecaller (Fig. 1a) [51, 52]. From the base probability matrix H, we extract the most probable basecalled sequence Y = {y*_τ_* } and its corresponding alignment A(t) which aligns the signal point time t to sequence index τ, giving t → τ . After correcting the basecalled sequence with the reference genome, we construct a reference k-mer sequence C by sliding a window of size k (in our case, k = 5) across the basecalled sequence, moving one base at a time. Each windowed segment forms a k-mer and is added to the sequence C = {c*_τ_* }. From now on, to simplify the notation, we use c*_t_* to denote the corresponding k-mer at time t after transitioning through alignment c*_A_*_(*t*)_. All time offsets of the k-mer sequence reside in the sequence domain, meaning c*_t_*_−1_ refers to c*_A_*_(*t*)−1_. Finally, we derived the k-mer transition matrix Ψ from k-mer sequence C; for details, see the next section. Then, the likelihood of observing an electrical signal X is given by:

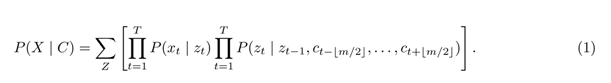

Here, *Z* is the hidden state representing the underlying k-mer sequence, z*_t_* is the k-mer at time t, and c*_A_*_(*t*)_ is the corrected k-mer representation at time t acquired from the canonical neural network output H (Fig. 1a). T is the maximum time stamp for a given sequence segment. m is the window size for the k-mers to be considered. P (x | z) is the emission probability of the signal x given the k-mer z, as modeled by a Gaussian distribution.

### Constructing a transition matrix from the base-called sequence and its alignment

We loosely constrain the transition matrix at time t in the nonhomogeneous HMM by using the base prediction output H derived from a canonical basecaller, thereby using the segmentation results provided by the basecaller in an error-tolerant manner (Fig. 1b). By calculating the most probable path from H, we can obtain both the basecalled sequence and the alignment between each base within the most probable path and the sequencing time t. Following this, we correct the basecalled sequence using the reference genome, and we also make appropriate revisions to the alignment to address the deletion or insertion errors in the basecalled sequence. We transform the corrected sequence into a k-mer sequence C = {c*_t_* : t = 1, . . ., T }, incorporating the k bases surrounding each base in the basecalled sequence; then, this k-mer sequence is reformatted into transition matrices Ψ = {ψ*_t_* : t = 1, . . ., T } by including at most m transitions, where each ψ*_t_* is the temporal transition matrix at time t. During the process of constructing the k-mer sequence C from H, the basecalled RNA sequence is corrected by aligning it to a reference genome through the following steps:

**–** For mismatched bases, we replace the bases in the k-mer with the reference bases.
**–** For insertions/deletions in the base-called sequences that are smaller than five bases, we determine the new signal alignment boundary of the inserted/deleted bases by evenly merging/splitting the signal boundaries of nearby bases; i.e., we redistribute the occupancy of the inserted bases to the nearby bases and allocate occupancy for the deleted bases from the nearby bases.
**–** We skip the sequence segments with insertions and deletions that are larger than five bases for quality control purposes.

The transition matrix Ψ is then constrained by C, masking out the irrelevant transition paths so that only transition paths that are likely to occur at time t are retained. To more clearly see what these temporal transition matrices stand for, let ψ*^t^* = Pr(z*_t_* = i | z*_t_*_−1_ = j, c*_t_*_−⌊*m/*2⌋_, . . ., c*_t_*_+⌊*m/*2⌋_) be the transition probability from k-mer i to k-mer j given constraint k-mers c*_i_* from a time window with a width of at most m, i.e., from t − ⌊m/2⌋ to t + ⌊m/2⌋. At the start and end of sequence, the window size is less than k due to boundary constraints. In comparison with the transition matrix ϕ*_i,j_* = P (z*_t_* = i | z*_t_*_−1_ = j) of a homogeneous HMM, the transition matrix now changes over time t:

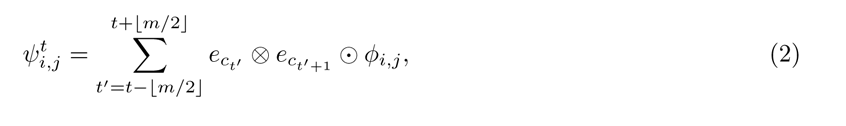

where ⊗ is the tensor product operation, ⊙ denotes elementwise multiplication, e*_i_*is a one-hot vector where only the i*^th^* element is 1, and ϕ*_i,j_* is the transition matrix in which ϕ*_i,j_*= 1 if the transition from k-mer i to k-mer j is valid (otherwise, it is 0). For example, AAACT to AACTA is valid, while AAACT to ACTCC is not, as we only allow 1 base step. ψ*^t^*is the k-mer transition matrix from the k-mer sequence described above; it is a binary value matrix indicating the k-mer transition i → j at time t, where 1 denotes a possible transition and 0 represents an impossible transition.

We construct the transition matrix from m nearby k-mers instead of only the k-mer at time t from kmer sequence C because the base probability predicted by the canonical basecaller is not exact due to the connectionist temporal classification (CTC) loss used [51, 52] and the insertion/deletion errors in the sequence, nor is it totally correct due to the previously unseen modified bases. Thus, we allow the NHMM to explore the alignment space in two ways. First, at each time point, the transition matrix of the NHMM is restricted to the current transition probability and the m nearby transition probabilities, where m is a hyperparameter (Eq. 2). This is done to make sure that the final alignment output by the NHMM is not too far away from the given the alignment from canonical basecalling but still allows for exploration within the m-base window. Second, the transition path of the underlying Markov chain is broadened to encompass all possible modified counterparts for each k-mer along the path (Fig. 1c). As an example, AACGT is extended to include four alternative k-mers with modified bases, AACGT (the original k-mer), AMCGT, MACGT, and MMCGT, leading to expanded paths. After the transition matrix is constructed for all the time points, the NHMM is then trained using the expectation-maximization (EM) algorithm [53] until it converges (Fig. 2b).

### Preparing the training data with data augmentation and read sampling

All-or-none methylated reads exhibit either complete methylation of all adenine (A) bases or none at all, whereas in actual biological samples, methylation typically occurs less frequently and is more sporadically distributed. To prevent the neural network from overfitting to all-or-none methylation reads, we create a training dataset containing partially methylated reads with labels. This is accomplished by dividing the signals from the all-or-none modified reads into smaller segments and subsequently splicing them together. The corresponding sequences are recombined according to their alignment with the signal, as provided by the NHMM. Merging the signals generated from distinct k-mers at their junction points can result in substantial discrepancies between the combined signal and the actual signal obtained from a real sequencing run. To avoid such deviations caused by k-mer mismatches, we ensure that the preceding and succeeding k-mers at the joint sections are identical. For instance, we can merge the signal segments with base-called sequences such as GGM***CGTTC*** XXX and XXX***CGTTC*** TAG to form GGM***CGTTC*** TAG. To achieve this, we define nonmethylatable k-mers as k-mers without adenine (***CGTTC*** in the example). They have the same sequencing signal distributions in both modified and unmodified reads, making them suitable for use as joint anchors. We employ the trained NHMM to decode both the canonical and fully modified reads in the training IVT dataset, using the base probability prediction from the canonical basecaller as described before. The alignment between the sequence and signal is established through a Viterbi path, which assigns each signal point to its corresponding k-mer (Fig. 1d). Each read is subsequently divided into segments at nonmethylatable k-mers. These segments are used to construct a k-mer signal graph, where each node represents an invariant k-mer. Each edge corresponds to a signal segment whose aligned sequence begins and ends at the respective k-mers of the connected nodes (Fig. 1e). We then perform a random walk on the graph, choosing the next edge via an ɛ-greedy sampling strategy with an upper confidence bound (UCB) [54], as used in the multi-armed bandit algorithm, to ensure maximum diversity in the sampling sequence (see Algorithm 1.4 in the supplementary materials).

### Data processing

#### Acquisition and processing of direct RNA sequencing datasets

All datasets used in this study are acquired from refs [28, 31, 36, 55]. We obtained both replicates (replicate 1 and 2) from the Epinano synthesized IVT RNA dataset [31] and the only single replicate from the ELIGOS synthesized IVT RNA dataset [28]. Both of these datasets contain fully modified reads and unmodified control reads. We also obtained all the NA12878 IVT RNA reads from the Oxford Nanopore human reference dataset repository: https://github.com/nanopore-wgs-consortium/NA12878/blob/master/RNA.md [55]. For the yeast dataset, we obtained all three replicates of the wild strain and *ime4*-knockout strain (ime4Δ) [31]. Reads are extracted if mapped to m6A-modified RRACH sites previously identified by antibody immunoprecipitation [37]. For the human HEK293T cell line, we obtained two replicates (replicate 1 and 2) of the wild-type human HEK293T cell [36] to evaluate models. Following a previous study [36], we used the reference transcriptome and its genome annotation provided by SG-NEx project: https://github.com/GoekeLab/sg-nex-data [41]. We used the same m6A DRACH sites in the m6Anet paper [36], which were originally identified by m6A-seq and miCLIP experiments [9, 12]. All replicates in the datasets are biological replicates, which are independent biological samples sequenced using the same direct RNA nanopore sequencing protocol. As for synthesized IVT reads, RNA replicates were transcribed from synthesized DNA reads with different sequences. See the sections below for details on replicates used for training and evaluating. All samples were generated using the Nanopore R9.4.1 flow cell, except for the human IVT data, which came from the R9.4 flow cell. The only significant difference between the two flow cells is the slightly improved yield in the R9.4.1.

#### Canonical basecalling and mapping

All reads in the training dataset were basecalled using the Guppy 5.0.11 ONT basecaller [56] and then mapped to the reference genome using minimap2 v2.24 [57] with the settings “-ax map-ont -uf --secondary=no --MD”. The mapped reads were then transferred to the BAM format using Samtools 1.11.0. A canonical neural network basecaller with the same structure as the CRNN was then trained using the NA12878 IVT reads, and this basecaller was then used to produce the base probability prediction. This canonical basecaller is used as a starting model when we retrain it on the augmented IVT data and subsequently fine-tune it on the yeast data [31].

#### Training datasets

We randomly selected 300,000 canonical (unmodified) read chunks and 300,000 fullymodified read chunks from replicate 1 of each of the two synthesized IVT RNA datasets [28, 31], as well as the first 300,000 canonical read chunks from the Oxford Nanopore Human IVT reference dataset [55] to construct the k-mer signal graph we described above. Reads were filtered out if the corresponding basecalled sequence was shorter than three bases, if the signal had a dwell time (the putative duration a k-mer remains in the pore) exceeding 2000 signal time points, if the basecalled sequence could not be aligned to the reference genome, or if a single base type comprised more than 60% of the basecalled sequence. This filtering process resulted in 228,983 canonical read chunks and 204,822 methylated read chunks from the first synthesized IVT dataset [31], 195,161 canonical read chunks and 213,085 methylated read chunks from the second synthesized IVT dataset [28], and 188,004 canonical read chunks from the Human IVT reference dataset [55]. Methylation sites identified by antibody immunoprecipitation [37], derived from the first replicate of the wild-type and the first replicate of the ime4Δ from the yeast dataset [31] were used to create the fine-tuning dataset. We regarded all sites from the wild-type strain as methylated and all sites from the ime4Δ strain as unmethylated. However, we considered these classifications noisy labels and used label smoothing during fine-tuning. Human HEK293T cell dataset [36] was not used for training and only used in the evaluation.

#### Evaluation datasets

All the accuracy evaluation datasets we used are sourced from previously published resources. These include a synthesized IVT dataset [31], a yeast dataset [31], and a human HEK293T cell dataset [36]. We used the second replicate from both the synthesized IVT and yeast datasets, as we had already used the first replicate of these two datasets for training and fine-tuning, and we used the first replicate of the human HEK293T cell dataset as it was not included in training. A subset of the human HEK293T cell dataset containing 500 genes was randomly sampled from the original dataset. For the yeast data, we assessed model performance based on the sites identified by m6A-seq [37] for the wild-type strain, and the ime4Δ strains where no methylation should be observed. For evaluation on human data, following previous work [36], we regarded the combined sites identified by m6A-seq [14] and miCLIP [12] as methylated sites, and other randomly selected sites with the DRACH motif as unmethylated sites.

### Training and fine-tuning a m6A methylation-sensitive neural network basecaller

We used the partially modified reads sampled from the signal k-mer graph to retrain a canonical basecaller. Before performing retraining on the pre-trained canonical basecaller, we reinitialized the parameters of the last fully connected hidden layer with random weights but kept the same standard deviation. We then retrained the model using a smaller learning rate (0.00001) than the usual learning rate (0.001). We finetuned our model on biological samples with m6A sites identified by antibody experiments [31], labeling the A base at each modified site as an m6A base for every read (Fig. 2b). Since the bases at methylation sites are usually not methylated in every read, this approach would introduce many false-positive labels. To address this issue, we applied label-smoothing to the connectionist temporal classification (CTC) loss that was used to train the basecaller. A label sequence of length L was defined as S = {s*_i_* : i = 1, 2, . . ., L}, and each s*_i_*belonged to the set {A, C, G, T, M}. The base probability logit output H ∈ R*^T/K^*^×*N*^ was a (T/K)-by-N matrix derived from the basecaller’s CRNN, where K is the total number of strides (i.e., the number of steps the convolutional filter moves across the input at each operation), and N is the number of bases used for prediction plus 1 (a blank symbol). The altered CTC loss with label smoothing under a strength factor represented by ɛ was then defined as:

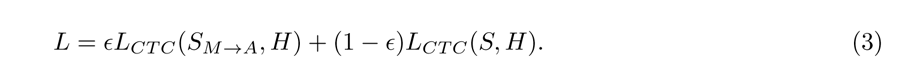

where M stands for the m6A base, L*_CT C_* is the usual CTC loss, and S*_M_*_→*A*_ is the sequence in which every m6A base is replaced with an A base. We set ɛ = 0.1 empirically for the fine-tuning process, with an expectation that the methylation label is correct with probability 1 − ɛ.

## 1 Supplementary Text

### 1.1 Summary of prior approaches

**Table S1.** Prior approaches concerning the identification of m6A modifications. AON: All-or-none modified dataset. KO: Knockout dataset.

| Method | Structure | Training data | Input features |
| --- | --- | --- | --- |
| Epinano (2019) [31] | SVM | AON | Basecall error & Signal feature |
| ELIGOS (2021) [28] | Stat. anal. | – | Basecall error |
| Nanocompore (2021) [29] | GMM | – | Signal features |
| nanom6A (2021) [33] | GBM | AON | Signal segments |
| CHEUI (2022) [35] | CNN | AON | Signal segments |
| m6Anet (2022) [36] | MLP | In vivo KO | Signal features & Sequence |
| Xron (This work) | CRNN | AON & In vivo KO | Raw signals |

### 1.2 K-mer encoded as integer

We encoded each k-mer with an integer by initially converting the k-mer string into a base-b integer. For example, ‘ACGTM’ is represented as a base-5 integer 01234_5_. This base-5 integer is then converted into a base-10 integer (z*_t_*), where 01234_5_ is transformed to 112_10_.

### 1.3 Signal segmentation

To determine the exact alignment between the raw current signals and the corresponding transcription positions, a signal segmentation procedure is typically required to assign consecutive signal points (called an event) to each base pair. The electrical current signals acquired from the ONT sequencer are 1D timeseries signals sampled at 4,000 points per second. Under the direct RNA sequencing protocol, the average movement speed of RNA through the pore is 70 base pairs per second, resulting in an average of 57 sampling points per base pair. The signal level and duration of an event are decided by the five nucleotides inside the pore, where the middle nucleotide is the one to which we mapped.

### 1.4 Sampling algorithm

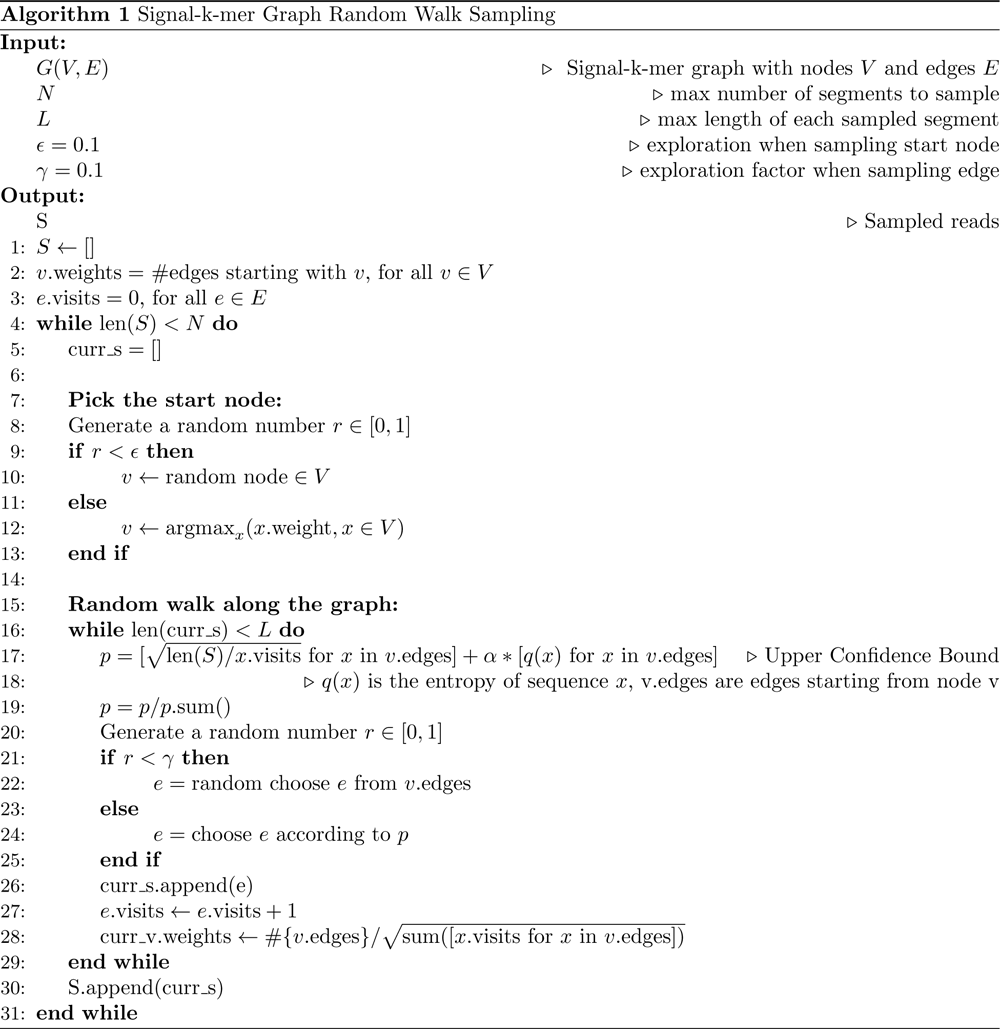

|  |  |
| --- | --- |
| $G(V, E)$ | ▷ Signal-k-mer graph with nodes $V$ and edges $E$ |
| $N$ | ▷ max number of segments to sample |
| $L$ | ▷ max length of each sampled segment |
| $\epsilon = 0.1$ | ▷ exploration when sampling start node |
| $\gamma = 0.1$ | ▷ exploration factor when sampling edge |
| $S$ | ▷ Sampled reads |

1: $S \leftarrow []$ 2: $v.weights = \#edges \text{ starting with } v, \text{ for all } v \in V$ 3: $e.visits = 0, \text{ for all } e \in E$ 4: while $\text{len}(S) < N$ do 5: $curr\_s = []$ 6: 7: Pick the start node: 8: Generate a random number $r \in [0, 1]$ 9: if $r < \epsilon$ then 10: $v \leftarrow \text{random node} \in V$ 11: else 12: $v \leftarrow \text{argmax}_x (x.weight, x \in V)$ 13: end if 14: 15: Random walk along the graph: 16: while $\text{len}(curr\_s) < L$ do 17: $p = [\sqrt{\text{len}(S)/x.visits} \text{ for } x \text{ in } v.edges] + \alpha * [q(x) \text{ for } x \text{ in } v.edges]$ ▷ Upper Confidence Bound 18: ▷ $q(x)$ is the entropy of sequence $x$ , $v.edges$ are edges starting from node $v$ 19: $p = p/p.sum()$ 20: Generate a random number $r \in [0, 1]$ 21: if $r < \gamma$ then 22: $e = \text{random choose } e \text{ from } v.edges$ 23: else 24: $e = \text{choose } e \text{ according to } p$ 25: end if 26: $curr\_s.append(e)$ 27: $e.visits \leftarrow e.visits + 1$ 28: $curr\_v.weights \leftarrow \#\{v.edges\} / \sqrt{\text{sum}([x.visits \text{ for } x \text{ in } v.edges])}$ 29: end while 30: $S.append(curr\_s)$ 31: end while

**Table S2.** Basecalling accuracy comparison between Xron and Guppy on three different datasets and their control datasets. The deletion, insertion, and mismatch rates (%) were calculated as the numbers of deleted, inserted, and mismatched bases divided by the number of bases in the reference sequence, respectively. The identity rate (%) was defined as the number of matched bases in the query sequence divided by the number of bases in the reference sequence (the higher the better). All reported rates are mean values among the aligned reads.

| Condition | Model | Deletion rate (%) | Insertion rate (%) | Mismatch rate (%) | Identity rate (%) (↑) |
| --- | --- | --- | --- | --- | --- |
| IVT Control | Xron | 4.14 | 11.60 | 8.51 | 87.35 |
|  | Guppy | 4.30 | 2.20 | 2.95 | 92.75 |
| IVT m6A | Xron | 5.09 | 15.04 | 6.44 | 88.48 |
|  | Guppy | 9.11 | 4.45 | 12.60 | 78.28 |
| Yeast ime4Δ KO | Xron | 9.47 | 4.54 | 5.57 | 84.97 |
|  | Guppy | 4.97 | 2.80 | 2.54 | 92.50 |
| Yeast | Xron | 9.12 | 3.83 | 6.92 | 83.96 |
|  | Guppy | 4.80 | 2.38 | 3.26 | 91.94 |
| HEK293T Mettl3 KO | Xron | 10.41 | 1.91 | 3.68 | 85.91 |
|  | Guppy | 4.42 | 2.59 | 2.39 | 93.19 |
| HEK293T | Xron | 9.46 | 2.08 | 3.43 | 87.12 |
|  | Guppy | 11.31 | 2.45 | 3.05 | 85.64 |

**Supplementary Figure 1.**
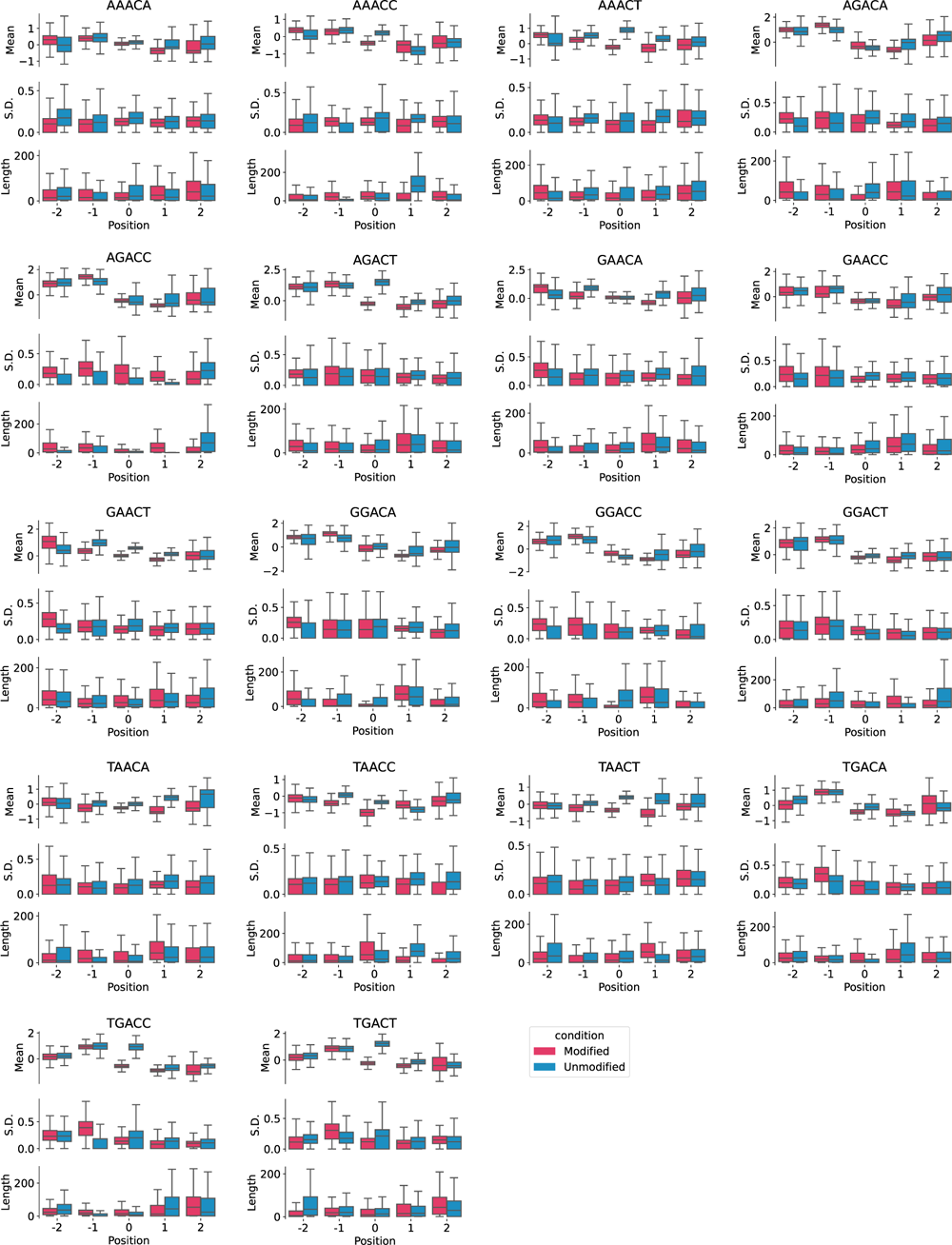
Signal features comparison for all DRACH motifs between modified and unmodified sites extracted from the Epinano IVT dataset. Box plot comparing the distributions of mean, standard deviation, and length between modified and unmodified sites for all 18 DRACH motifs. Horizontal lines show the median, the box denotes the interquartile range, and the whiskers extend to 1.5 times the interquartile range. Points beyond this range are considered outliers and are removed from the plot.

**Supplementary Figure 2.**
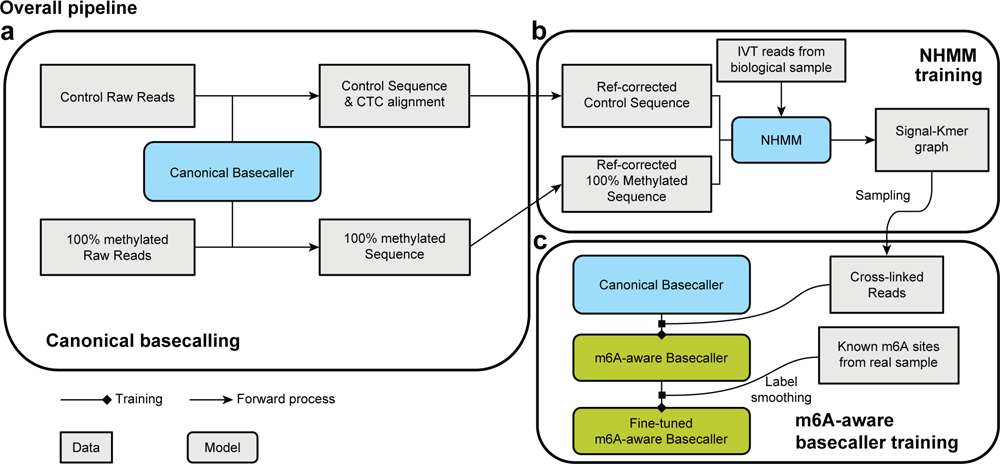
Overall training pipeline of Xron training. **a** Basecalling the modified and un-modified reads using a canonical basecaller. **b** Training the NHMM with the corrected synthesized RNA sequence and IVT reads from human reference data. The trained NHMM was used to generate a signal k-mer graph. **c** The Xron m6A-distinguishing Basecaller was trained using the cross-linked reads sampled from the signal k-mer graph and then fine-tuned on the yeast and human datasets, where putative m6A sites were identified through an immuno-precipitation experiment. We applied label smoothing when fine-tuning the model due to the noisy m6A labels, as the m6A modification for each read was unknown.

**Supplementary Figure 3.**
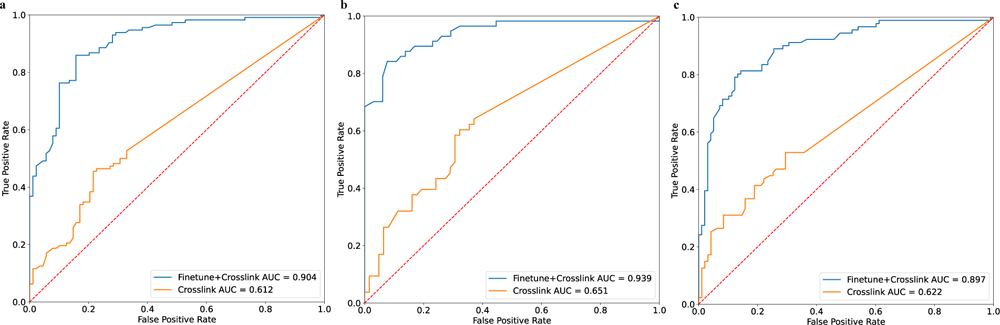
Ablation study of Xron model. To validate the necessity of finetuning Xron on IP data, an ablation study was conducted. We evaluate the performance of Xron on three biological replicates (a-c) of yeast data, with and without IP data finetuning. The plots show a dramatic decrease in model performance without finetuning using IP data. Xron model was finetuned using the first replicate of the yeast data.

